# RACK1 may participate in placental development via regulating proliferation and migration of trophoblast cell in pigs following intrauterine growth restriction

**DOI:** 10.1101/2022.09.29.510071

**Authors:** Zhimin Wu, Guangling Hu, Ting Gong, Qun Hu, Linjun Hong, Yiyu Zhang, Zheng Ao

## Abstract

Intrauterine growth restriction (IUGR) is a severe complication in swine production. Placental insufficiency is responsible for inadequate fetal growth, but the specific etiology of placental dysfunction-induced IUGR in pigs remains poorly understood. In this work, placenta samples supplying the lightest-weight (LW) and mean-weight (MW) pig fetuses in the litter at day 65 (D65) of gestation were collected, and the relationship between fetal growth and placental morphologies and functions was investigated using histomorphological analysis, RNA sequencing, quantitative polymerase chain reaction, and in-vitro experiment in LW and MW placentas. Results showed that the folded structure of the epithelial bilayer of LW placentas followed a poor and incomplete development compared with that of MW placentas. A total of 632 differentially expressed genes (DEGs) were screened out between the LW and MW placentas, and RACK1 was found to be downregulated in LW placentas. The DEGs were mainly enriched in translation, ribosome, protein synthesis, and mTOR signaling pathway according to GO and KEGG enrichment analyses. In-vitro experiments indicated that the decreased RACK1 in LW placentas may be involved in abnormal development of placental folds (PFs) by inhibiting the proliferation and migration of porcine trophoblast cells. Taken together, these results revealed that RACK1 may be a vital regulator in the development of PFs via regulating trophoblast ribosome function, proliferation, and migration in pigs.

## 1 Introduction

Intrauterine growth restriction (IUGR) is a common obstetrical complication resulting in many adverse effects on fetal growth and development and postnatal health [1, 2]. Pigs are one of multiparous species most likely to suffer from IUGR among domestic animals (accounting for 15%–25% of births), though a good deal of measures has been made to reduce the occurrence of IUGR, which negatively influences the production performance and economic benefits of pig production [3]. Piglets with IUGR have been shown to be followed by high morbidity and mortality, and they are predisposed to stunted growth, digestive diseases, and poor carcass quality [4-6]. Although the etiology of IUGR derives from maternal, fetal, placental, or genetic causes, accumulating evidence in humans and experimental animals demonstrated that the majority of IUGR cases principally point to a failure of the placenta associated with a decrease in maternal-fetal nutrients and oxygen exchange [1, 7, 8]. However, the pathophysiological mechanisms underlying placental dysfunction-induced IUGR in pigs remain poorly understood.

The placenta is a transient organ that only persists for the duration of pregnancy but is absolutely crucial for all intrauterine events [9], fulfilling key tasks to transporting nourishment, producing hormones and cytokines, acting as a waste filtration system, and as a protective barrier to guarantee physiological adaptations of mother and fetus during pregnancy [10, 11]. Porcine placenta belongs to epitheliochorial type, where columnar trophoblasts lack significant invasion but spread loosely over the uterine luminal epithelial layer to form the folded bilayer [12]. Previous studies have indicated that the reduction in utero-placental blood flows and/or angiogenesis contribute to insufficient transport of nutrients and fetal hypoxia that is likely to be related to formation of IUGR fetuses [8, 13]. Interactions between placental trophoblast cells and maternal immune cells are also known to have an influence on the growth trajectories of a fetus [9]. Proteomics analysis revealed that the placenta and endometrium of IUGR pig fetuses are vulnerable to nutrient transport reduction, oxidative damage, and impairment of cell metabolism [14]. Placental structure is also an important factor in determining placental efficiency. Previous studies showed that the placental folds (PFs), especially the shape of trophoblast cells, and the expression of regulatory genes of Meishan pigs underwent more complex changes than those of Yorkshire pigs, and such changes may be a potential factor for their differences in reproductive performance.

The placenta possess a unique transcriptional landscape throughout pregnancy, so its growth and development are regulated by sophisticated pathways composed of the expression of substantial genes [11, 15]. Incorrect alterations in gene expression in placenta give rise to abnormal morphologies and dysfunction have been found to be associated with various pregnancy complications [11, 16]. The development of pig placenta reaches completion in terms of weight, surface area, and numbers of placental areolae by days 60–70 of gestation [17], which is crucial for fetal growth and development in late gestation. However, transcriptome analysis of placenta related with pig IUGR fetuses in this period is rarely reported. The present study aimed to identify the differentially expressed genes (DEGs) in the placentas of fetuses with lightest weight (LW) compared with those in the placentas of mean-weight (MW) litter during day 65 (D65) of gestation through mRNA sequencing, and illustrate the function of RACK1 on porcine trophoblast cells. The findings could provide basic reference for future etiological mechanisms of IUGR in pigs.

## 2 Materials and Methods

### 2.1 Ethics statement

All experimental design and protocols in this study were reviewed and approved by the Institutional Animal Care and Use Committee of Guizhou University, Guiyang, China (EAE-GZU-2020-T010). All efforts were made to minimize animal suffering.

### 2.2 Tissue collection

Five Duroc sows showing signs of spontaneous estrus and with similar litter size record, parity, and weight were artificially inseminated twice daily with the semen from the same Duroc boar. All sows were raised under similar conditions. After the pregnant sows were hysterectomized after the induction of anesthesia (xylazine, 2.0 mg/kg bw) during D65 of pregnancy, the uteri were opened from the corners. At the time of dissection, all fetuses were identified as “live” or “dead,” and their sex was determined on the basis of their morphology. Each fetus was weighed, and the corresponding placenta sample was collected and immediately snap-frozen in liquid nitrogen and stored at −80 °C. The LW fetuses and those closest to the MW of the litter were identified and chosen on the basis of fetal body weights. Meanwhile, the placenta samples were fixed in 4% paraformaldehyde used for histomorphological examination.

### 2.3 Histomorphological analysis

The placental tissues were fixed in 4% paraformaldehyde for a minimum of 48 h and then embedded in paraffin, sectioned with 5 μm thickness, and stained with hematoxylin–eosin (H&E) on the basis of standard histological criteria. Subsequently, placental histomorphometry of the stained sections was executed as described previously [18, 19]. Placental data were obtained using a Nikon Ni-U light microscope (100× magnification) fitted with a Nikon (DS-Fi1) digital camera (Nikon, Japan). Morphometric measurements of the average width of the PFs and fold length (μm) per micrometer of placenta were calculated using ImageJ 1.45 software (National Institutes of Health, Bethesda, MD).

### 2.4 Total RNA isolation and RNA sequencing

Eight placenta samples of the fetus with LW (male, n = 4) and closest to the MW (male, n = 4) of the litter were selected for RNA sequencing. The total RNA was extracted from the placental tissues with the Total RNA Kit (Omega, USA) in accordance with the manufacturer’s instructions. The RNA quality and amounts were evaluated using a 2100 Bioanalyzer (Agilent Technologies, Santa Clara, CA) and agarose gel electrophoresis. All extractions exhibiting an RNA integrity number > 7.0 and a 28S:18S ratio > 1.0 were used in the next experiments. mRNA sequencing was carried out on the Illumina Hiseq 2500 system (Illumina, San Diego, CA, USA) and 150 bp paired-end FASTQ read files were generated. Raw data were deposited in the NCBI Sequence Read Archive database under accession number PRJNA838349. They were filtrated to obtain clean reads by removing adaptor and low-quality reads. The clean reads were aligned with the Ensembl *Sus scrofa* reference genome (version:scrofa10.2.87) by using Hisat2 version 2.0.5 [20], followed by transcript assembly and differential transcript expression analysis with Cufflinks version 2.2.1. Gene expression was measured with fragments per kilobase million mapped reads (FPKM) by using Cufflinks version 2.2.1. DESeq2 was used to normalize the read counts and apply FPKM values to calculate the relative gene expression differences by adjusting *P* values via Benjamini and Hochberg’s method to control the false discovery rate (FDR) and fold change (FC). Differentially expressed genes (DEGs) were screened out when the adjusted p-value < 0.05 and |log_2_FC| ≥ 1. Gene ontology (GO) annotation and Kyoto Encyclopedia of Genes and Genomes (KEGG) pathway analysis were performed on all DEGs by using KOBAS 3.0 (http://kobas.cbi.pku.edu.cn), and the DEGs were considered significant at FDR < 0.05.

### 2.5 Construction of protein–protein interaction (PPI) network

The Search Tool for the Retrieval of Interacting Genes (STRING) database (https://string-db.org/) is utilized to identify the pairwise relationships of all DEGs by computational prediction methods. In this study, STRING was used to predict the interactions among proteins encoded by candidate genes, with cutoff for confidence scores of interactions > 0.4. Cytoscape software (https://cytoscape.org/) was applied to visualize the results of the PPI network. Molecular Complex Detection (MCODE), a Cytoscape plugin, was used to locate the hub genes of the PPI network with the number of nodes > 10.

### 2.6 Quantitative real-time PCR (qRT-PCR)

cDNA was generated from 1 μg of total RNA by using the PrimeScript RT Reagent Kit (Takara, Japan) in accordance with the manufacturer’s protocol as previously described [21]. Quantitative PCR was performed on QuantStudio 3 (Applied Biosystems, MA, USA) following the parameters recommended by the manufacturer. Each reaction mixture (10 μL) contained 1 μL of cDNA solution, 0.3 μL of 10 mM of each specific primer, 5 μL of SYBR Select Master Mix, and 3.4 μL of ddH_2_O. The PCR reactions were run as follows: initial denaturation at 95 °C for 2 min; 40 cycles of denaturation at 95 °C for 15 s; annealing at 60 °C for 15 s, with an extension at 72 °C for 1 min; and finally, a melting curve was drawn at 95 °C for 15 s, 60 °C for 1 min, and 95 °C for 15 s. **Table 1** shows the primer information of target genes. Each gene expression test was performed in triplicates. The specificity of the PCR reaction was confirmed through a single peak in the melting curve. Gene expression levels were normalized with β-actin to calculate the relative expression levels by using the 2^−ΔΔCt^ method.

**Table 1.**
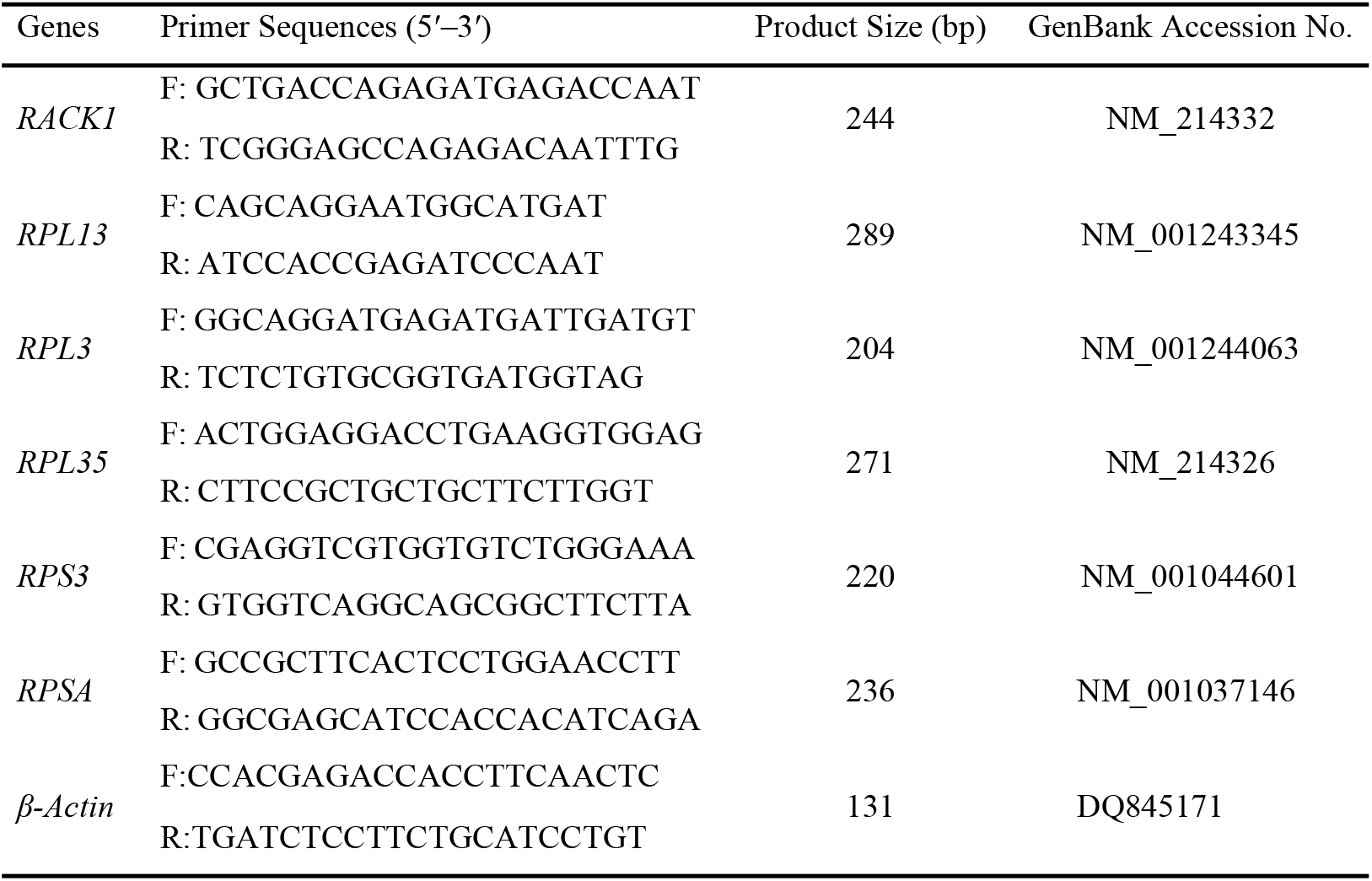
Primers used for qPCR.

### 2.7 Construction of RACK1 overexpression plasmid

The coding sequences (CDS) of the porcine RACK1 gene were amplified from the NCBI database (https://www.ncbi.nlm.nih.gov/nuccore/NM_214332.1). The RACK1 CDS was analyzed to identify appropriate restriction enzymes for vector construction using Primer 5.0 software (PRIMER-E Ltd., Plymouth, UK). The recombinant plasmid containing the RACK1 CDS and the pEGFP-C1 plasmid were subjected to digestion with *Sal*I and *Xba*I, and the linked product was subsequently used to transform competent DH5α. Endotoxin-free plasmids containing the correct fragment were identified by restriction enzyme digestion and sequencing.

### 2.8 Cell culture and transfection

Porcine trophoblast cell line PTr2 were kindly provided from South China Agricultural University as previously described [22]. The PTr2 cells were cultured in DMEM-F12 (Gibco, Waltham, MA, USA) supplemented with 10% fetal bovine serum (Gibco, Carlsbad, CA, USA) and recombinant human insulin (Yeasen, Shanghai, China) and maintained at 37 °C with 5% CO_2_ humidified atmosphere. They were then transferred into a six-well plate with 0.25–1 × 10^6^ cells per well and transfected with pEGFP-C1-RACK1 and pEGFP-C1 plasmid by Lipo8000 (Beyotime, Shanghai, China) in accordance with the manufacturer’s protocol.

### 2.9 Cell proliferation and migration assay

The cell proliferation in this study was assessed by cell counting kit-8 (CCK8) assay (Solarbio, Beijing, China). The PTr2 cells were seeded on 96-well plates with 100 μL of complete culture medium; treated with pEGFP-C1-RACK1, and pEGFP-C1 plasmid; and incubated for increasing durations (0, 12, 24, and 48 h). Then, 10 μL of the CCK8 solution was added per well and incubated for 2 h at 37 °C. Absorbance was measured at 450 nm with a microplate reader (CYTATION5, BioTek, USA).

Wound healing assay was performed to analyze the influence of overexpressed RACK1 on PTr cell migration as described elsewhere. In brief, 2 × 10^5^ cells were inoculated in F12 medium containing 10% FBS and 0.1% insulin. After the cells reached 80–90 % confluence, a perpendicular wound was established by creating a linear cell-free region with the use of a 10 μL pipette tip. The cells were washed with PBS twice, fresh complete medium was added, and the overexpressed vector and empty vector were transfected into the cells at the same time. The progress of cell migration into the scratch was photographed at 0 and 24 h after wounding with the use of computer-assisted microscopy. The images were quantitatively analyzed using ImageJ software (National Institutes of Health, Bethesda, MD). Cell migration was calculated as percentages of cell coverage to the initial cell-free zone.

### 2.10 Statistical analysis

The placental histomorphological data and mRNA expression levels were analyzed via one-way analysis of variance using SPSS 19.0 software (IBM Corp., Armonk, NY, USA). The data for the CCK8 and wound healing assays were evaluated using Student’s t-Test. All data are presented as mean ± standard error of mean (SEM). *P* < 0.05 was considered statistically significant.

## 3 Results

### 3.1 Fetal and placental characteristics in pigs at D65 of pregnancy

In this study, all five sows from artificial insemination were pregnant, and they produced 42 live and three dead fetuses at D65 of pregnancy. The average body weight of pig fetuses was 204.4 g, with a range of 134–259 g (**Fig 1A**). Corresponding maternal–fetal interface samples of fetuses with LW and MW of the litter were collected to investigate the associations of fetal weight and placental morphologies and functions. Next, the histomorphologies of the maternal–fetal interface of the LW and MW fetuses were compared by the sections stained with H&E. The results of observations revealed that the folded structure of the epithelial bilayer of the LW placentas followed a poor and incomplete development compared with that of the MW placentas (**Fig 1B**). The morphometry analysis showed that the fold width and fold length (μm) per micrometer of the LW placentas were extremely significantly lower than those of the MW placentas (**Fig 1C** and **1D**, *P* < 0.001).

**Fig 1.**
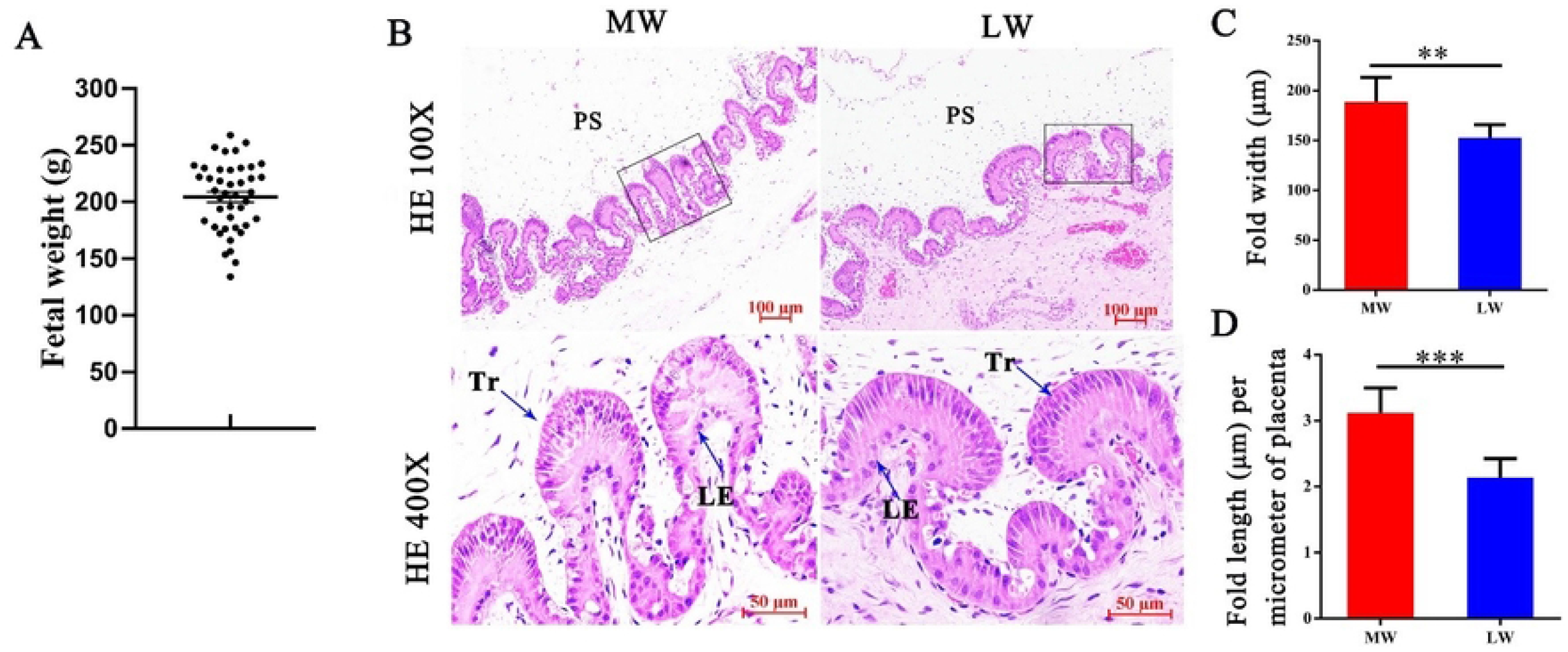
Characteristics of fetal weight and placental morphology in pigs during day 65 of gestation. (**A**) Distribution of fetal body weight. (**B**) Photomicrographs of representative sections of maternal–fetal interface derived from fetuses with lightest weight (LW) and litter with mean weight (MW) and stained with hematoxylin and eosin. Placental trophoblast (Tr) that came into touch with endometrial luminal epithelium (LE) to form placental folds (PF). (**C**) Width of PFs and (**D**) fold length (μm) per micrometer of placenta used to compare placental morphometry. PS, placental stroma; \*\**p* < 0.01; \*\*\**p* < 0.001

### 3.2 Differences in transcriptomic profiles between LW and MW placentas

After trimming for adapters and removing the low-quality reads, RNA-seq libraries that generated 40.6–41.7 million sequence reads among samples were used for downstream analysis. A total of 41,150,783 and 41,396,695 clean reads and 32,991,736 and 33,325,260 mapped reads were obtained from the LW and MW placentas, respectively (**S1 Table**). Through gene expression analysis (*q-*value < 0.05; FC > 2), 632 genes were identified to be significant DEGs between the LW and MW placentas (**S1 Fig**), and 535 and 119 genes were upregulated and downregulated in the LW placentas, respectively. Hierarchical clustering was performed with the datasets of DEGs. The mRNA expression patterns of the LW and MW placenta samples were clustered separately after clustering (**Fig 2**).

**Fig 2.**
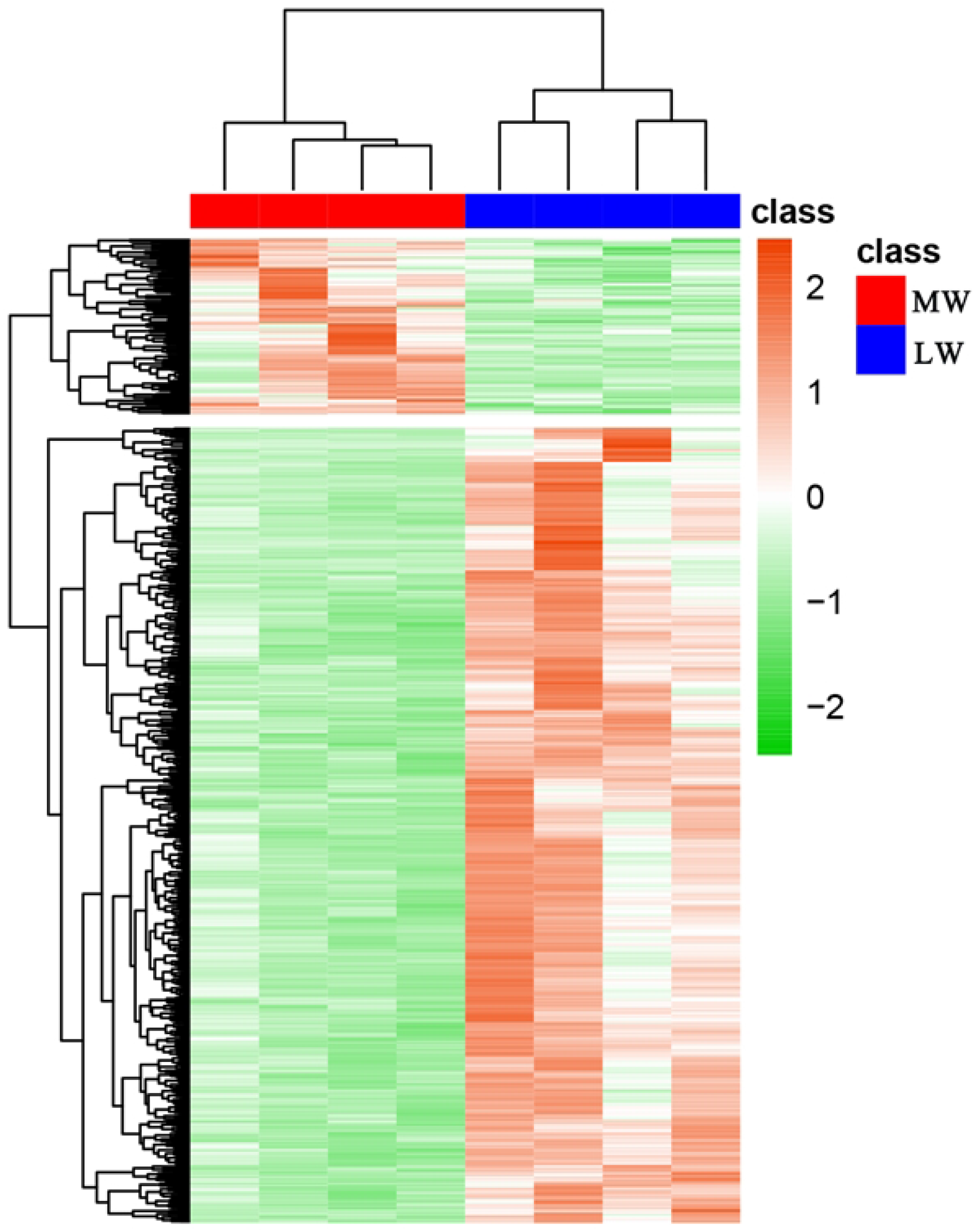
Heatmap of differentially expressed genes (DEGs) in LW and MW placentas. Orange and green represent genes with high and low expression levels, respectively.

### 3.3 GO function and KEGG pathway analysis of DEGs

GO enrichment analysis revealed that the DEGs mainly functionally enriched in biological processes were protein ubiquitination; signal transduction by protein phosphorylation; cytoplasmic translation; translational elongation; positive regulation of GTPase activity; positive regulation of transcription by RNA polymerase II; and negative regulation of transcription, DNA-templated (**Fig 3A**). The cellular components were mainly composed of polysomal ribosome, nucleus, cytosol, cytoplasm, and cytosolic large ribosomal subunit (**Fig 3B**). For GO molecular function, ATP binding, DNA binding, protein kinase binding, ribonucleoprotein complex binding, and RNA binding were the most significantly enriched terms. KEGG pathway enrichment analysis showed that the DEGs of the two groups of placentas participated in ribosome pathway, endocytosis, mTOR signaling pathway, phagosome, lysosome, and regulation of actin cytoskeleton.

**Fig 3.**
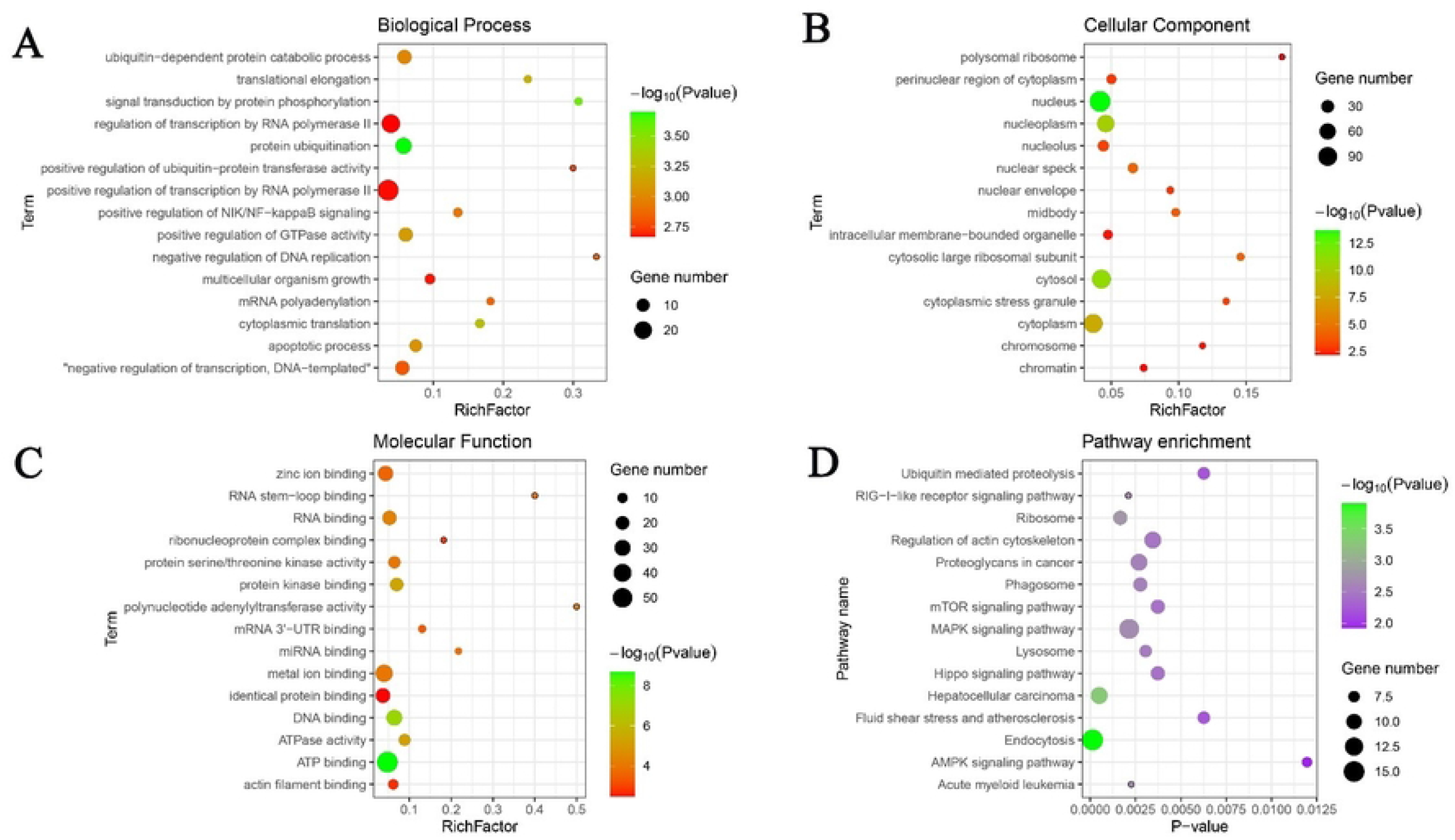
GO enrichment and KEGG pathway analysis of differentially expressed genes (DEGs) between LW and MW placentas. (**A**) Biological process term of GO enrichment analysis. (**B**) Cellular component term of GO enrichment analysis. (**C**) Molecular function term of GO enrichment analysis. (**D**) KEGG pathway analysis of DEGs. GO: Gene Ontology; KEGG: Kyoto Encyclopedia of Genes and Genomes.

### 3.4 PPI network and MCODE analysis

A PPI network of the DEGs with 544 nodes and 1263 edges was constructed through STRING analysis to identify the functions of the DEGs (**Fig 4A)**. The MCODE application in Cytoscape software was used to perform gene network clustering analysis to identify the key PPI network modules. The MCODE results with the parameter K-Core = 2 showed that one significant module was screened out from the PPI network (**Fig 4B**). A total of 15 nodes and 206 edges were found, and *RACK1, RPL13, EEF2, RPS3, ABCE1, EEF1D, RPL3, RPL10A, RPL35, RPLP1, RPL12, RPL18, RPL27A, EEF1G*, and *RPSA* were hub nodes in the module with score = 14.714 (**Fig 4B**). KEGG pathway analysis demonstrated that most hub genes were involved in the ribosome pathway based on KOBAS database (**Table 2**). Besides, RNA-sequencing results were confirmed by qRT-PCR. The results showed that in the LW placentas, the mRNA expression levels of *RACK1, RPL13, RPS3, RPL3*, and *RPL35* were significantly lower (*p* < 0.05) and *RPSA* did not differ (*p* > 0.05) compared with those in MW placentas (**Fig 5**). Increasing evidence have shown that RACK1 had roles on and off the ribosome (23, 24), suggesting that RACK1 may affect the growth and development of trophoblast cells via regulating ribosome function.

**Fig 4.**
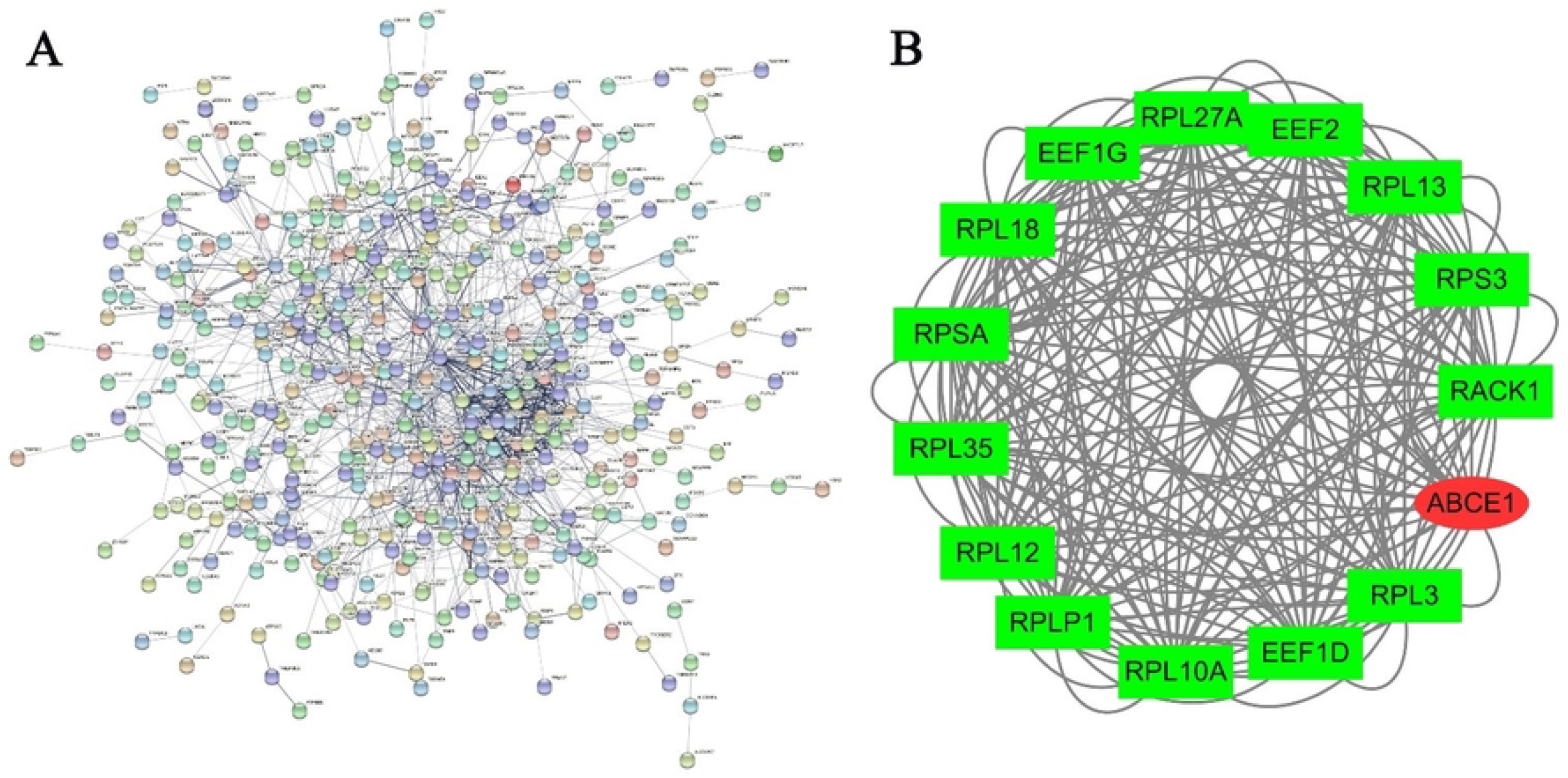
Protein–protein interaction (PPI) network and MCODE analysis. (**A**) PPI network constructed by STRING. The size of each node is positively correlated to the number of degrees. Interactions are shown by edges, with thicker edges corresponding to stronger associations. (**B**) MCODE analysis of differentially expressed genes. The red round nodes represent upregulated genes; the green square nodes represent downregulated genes.

**Table 2.**
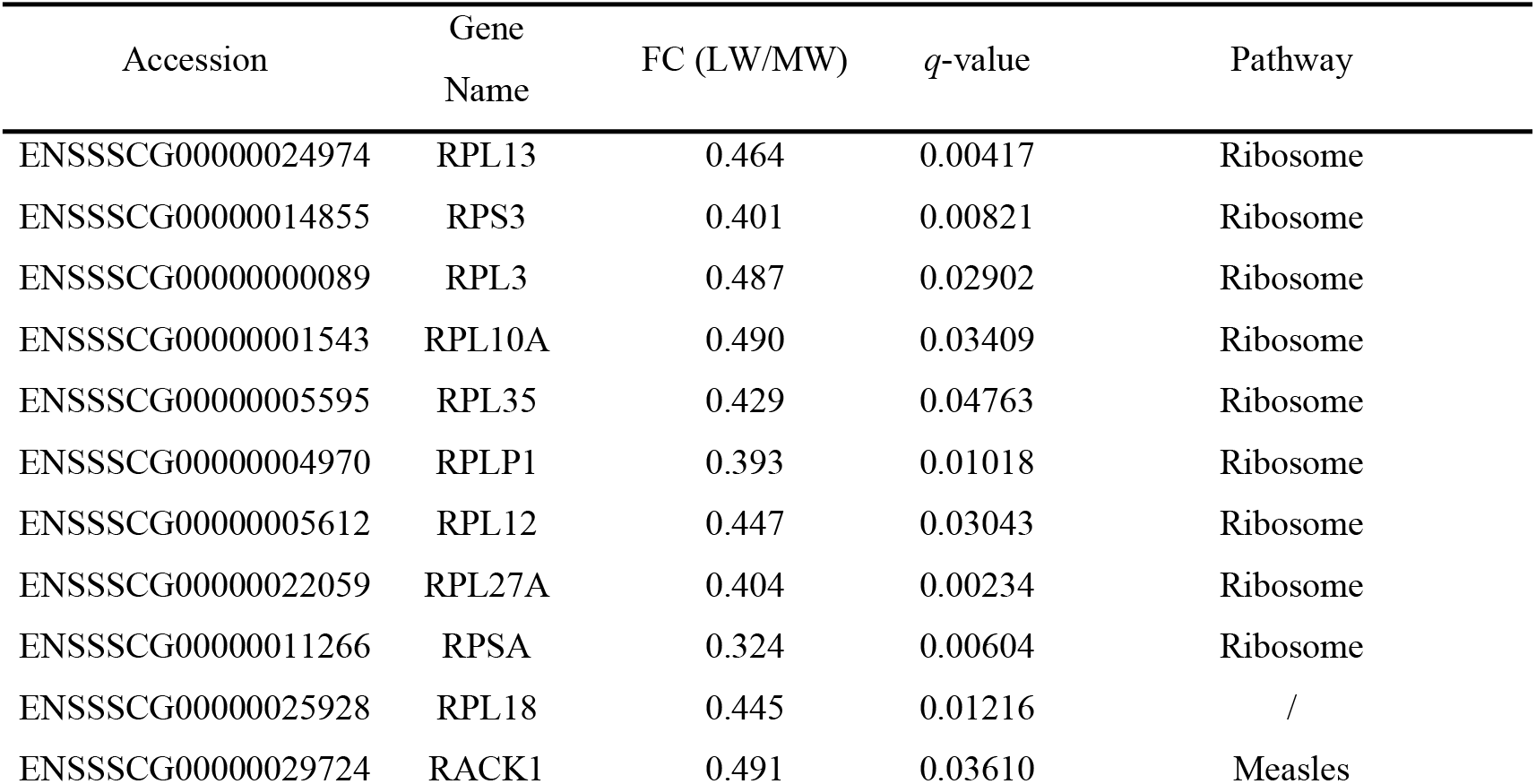
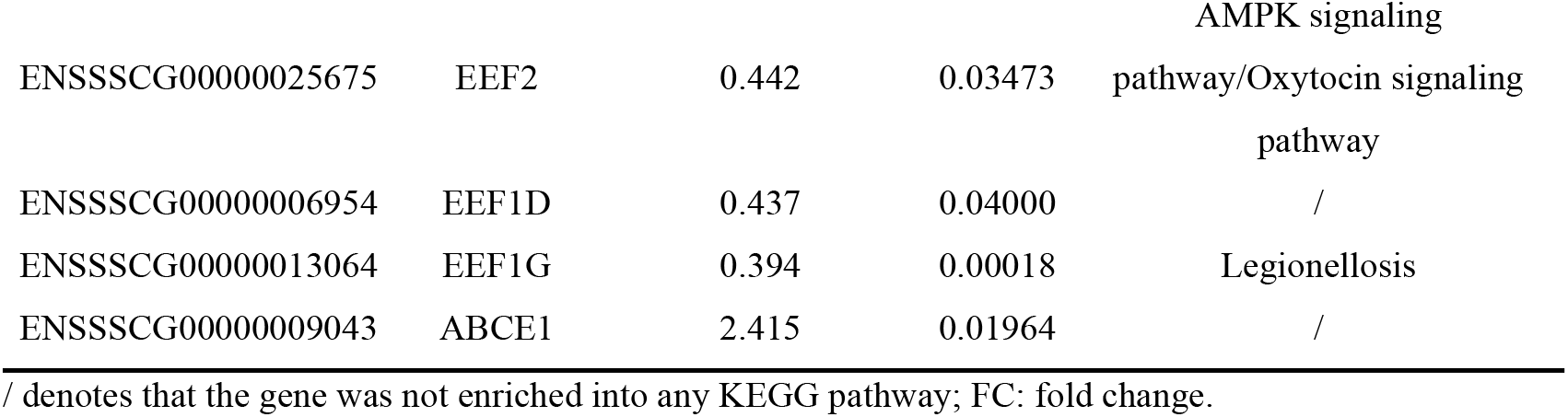
List of differentially expressed genes in a significant MCODE module from PPI network.

**Fig 5.**
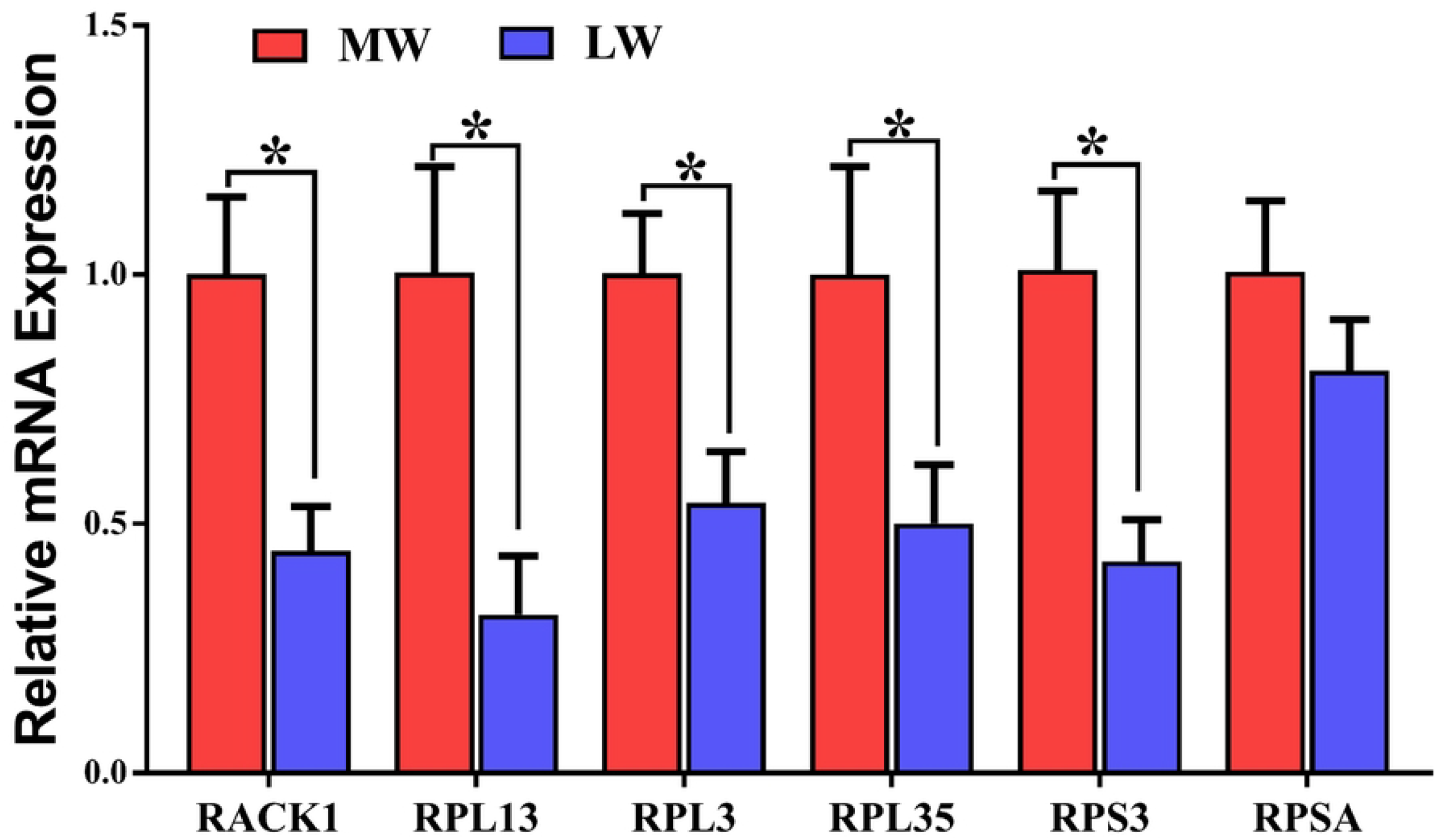
Validation of differentially expressed genes between LW and MW placentas by q-PCR. Values are presented as mean ± SEM. Values labeled with asterisk (*) indicate they are significantly different (*p* < 0.05).

### 3.5 Promotion of the proliferation and migration of PTr2 cells by RACK1 overexpression

RACK1 was overexpressed in PTr2 cells by transfection with pEGFP-C1-RACK1 plasmid to further explore whether it was involved in the biological behavior of trophoblast cells in this experiment (**S2 Fig**). The CCK8 assay revealed that overexpression of RACK1 significantly increased the proliferation of PTr2 cells compared to the corresponding negative control (**Fig 6**). Wound healing assays were also conducted To further confirm the role of RACK1 in trophoblast migration. The results demonstrated that overexpression of RACK1 increased the migratory ability of PTr2 cells compared with the corresponding negative control (**Fig 7**). These results suggested that RACK1 is required in the migration and proliferation of porcine trophoblasts.

**Fig 6.**
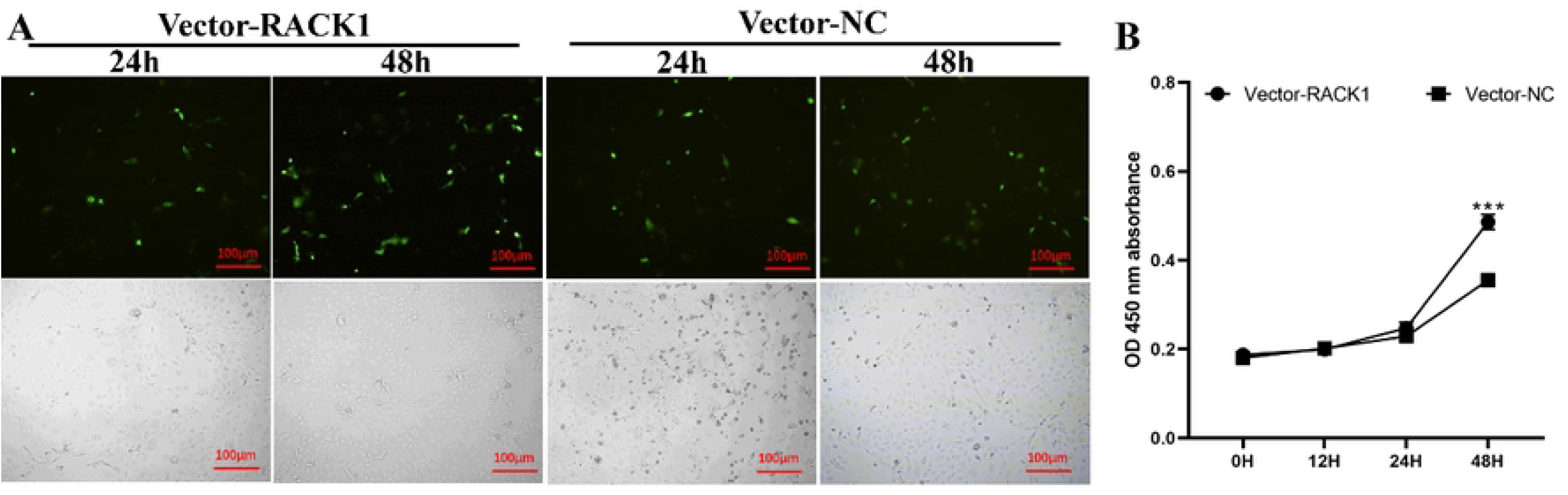
Effects of RACK1 overexpression on proliferation of PTr2 cells. (**A**) Microscopic results of PTr2 cells transfected with pEGFP-C1-RACK1 plasmid vector (vector-RACK1) and corresponding negative control vector (vector-NC) after 24 and 48 h. (**B**) Time-dependent cell viability of PTr2 cells with stable RACK1 overexpression measured by CCK-8 assay, which was evaluated using Student’s *t*-test. CCK8, Cell Counting Kit-8. Data are shown as mean ± SEM. \*\*\**P* < 0.001.

**Fig 7.**
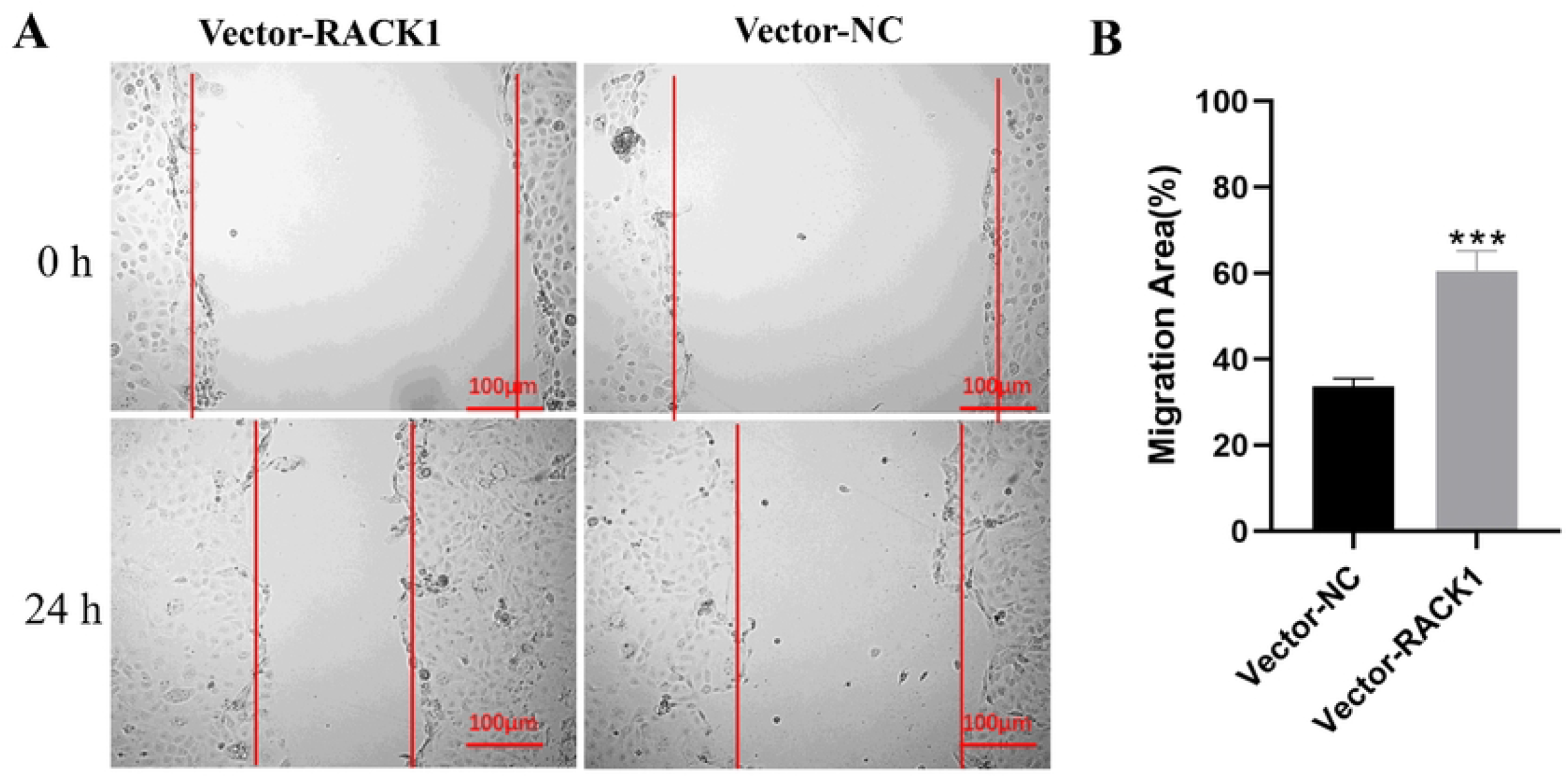
Effects of RACK1 overexpression on migration of PTr2 cells. (**A**) Representative wound healing photomicrographs and (**B**) quantitative analysis of wound healing rates of PTr2 cells with stable RACK1 overexpression transfection after 0 and 24 h. These tests were repeated three times independently. The wound healing assay was evaluated using Student’s *t*-test. Data are presented as mean ± SEM. \*\*\**P* < 0.001.

## 4 Discussion

Numerous studies during the past decades have confirmed that placental insufficiency is responsible for the abnormal growth trajectory of a fetus [25, 26]. Improved understanding of the mechanisms governing fetal growth is conducive to reduce the prevalence of low birthweight in pig production industry. Placental structure is compactly associated with its function and fetal growth and development. The results of the present study showed that low-weight pig fetuses had poor and incomplete PFs. In the epitheliochorial placenta, PFs could increase the contact area between trophoderm and endometrial epithelium to ensure that the fetus receives adequate nutrition from the maternal circulation. This finding was consistent with several previous findings that significant differences in placental microscopic folds between large and small pig fetuses due to the development of PFs have a substantial effect on placental efficiency [19, 27].

The development of PFs is associated with facilitating placental efficiency, so the efficiency of nutrient transport from the pregnant sow to the developing fetus depends on the size and function of the placenta [19]. Therefore, poor placental development may contribute to compromised nutrient transport that gives rise to low fetal body weight. The epitheliochorial placenta initially appears around days 26–30 of gestation, and regular PFs are formed on day 50 of gestation in pigs [22,28,29]. Recent findings in pigs revealed that large fetuses had higher trophoblastic epithelium of the chorioallantois fold than small fetuses at days 45 and 60 of gestation [30]. However, the width of the folded bilayer in the placentas of small fetuses was greater than that in the placentas of large fetuses on day 105 of gestation because PFs underwent rapid increase from mid- to late pregnancy that serves as a compensation in morphometric changes to increasing the surface area of interaction and then improving placental efficiency in response to low fetal weight and reduced placental size [13, 19].

RNA sequencing showed the differences in gene expression pattern between the LW and MW placentas and the multiple genes involved in the ribosomal pathway and mTOR signaling pathway in the LW placentas. Any disturbance in the ribosome environment could have devastating effects on placental development [31, 32]. Ribosomal proteins (RPs) are essential in regulating translation for facilitating placental growth and development [33]. Previous studies have found that many genes encoding ribosomal proteins were in the IUGR placenta [34]. mTOR signaling pathway have been reported to participate in the regulation of cellular activities of trophoblast in multiple pregnancy complications, including IUGR [35]. In human IUGR placenta, the downregulation of the expression of ribosomal proteins (RPL26 and RPS10) was regulated by the mTORC1 signaling pathway, affecting protein synthesis and leads to placental dysfunction [36]. Several reports have indicated that decreased protein synthesis and/or increased protein degradation is a constant feature during fetal growth restriction [37-39]. Hence, dysregulated expression of RP genes may alter the morphologies of LW placenta via affecting the translation and protein synthesis of trophoblast cells.

Through transcriptomic analysis and in-vitro experiment, this study further found that RACK1 potentially plays a critical role in placental development by regulating the proliferation and migration of porcine trophoblast cells. The migration of trophoblasts at the maternal-fetal interface after implantation is a critical process in placentation to facilitate the establishment of feto-placental circulation and thus essential for successful pregnancy outcomes in mammals [40]. Accumulating evidence has shown that RACK1 is involved in diverse biological processes, including protein translation, cell growth, cell cycle progression, cell migration, and stress responses, by localizing to different subcellular structures, including the nucleus, ribosome, and midbody [41-44]. This finding is consistent with the results of enrichment analysis of DEGs between the MW and LW placentas in the present study. Furthermore, studies on model animals have confirmed that knockdown of RACK1 suppresses cell growth in cultured cells and homozygous knockout of RACK1 is lethal for the embryo [45-47]. A recent study on pigs has shown that the proliferation and invasion of trophoblast cells affect the formation and development of PFs [27]. Therefore, the downregulation of RACK1 in the LW placentas may lead to abnormal development of PFs through inhibiting the proliferation and migration of porcine trophoblast cells, which possibly is a potential cause of fetal growth restriction.

In conclusion, this study demonstrated that the folded structure of the epithelial bilayer of placentas supplying LW fetuses followed a poor and incomplete development compared with that of the placentas supplying MW fetuses at D65 of gestation. A total of 632 DEGs were screened out between the LW and MW placentas, and they were mainly enriched in translation, ribosome, protein synthesis, and mTOR signaling pathway. The data also revealed that RACK1 was downregulated in the LW placenta. The decreased RACK1 in the LW placentas may be involved in the abnormal development of PFs by inhibiting the proliferation and migration of porcine trophoblast cells. Further works are required to elucidate the detailed molecular mechanisms underlying RACK1 in the regulation of porcine trophoblast cells.

## Author contributions

Zheng Ao and Zhimin Wu conceived and designed the study; Guangling Hu and Zhimin Wu and Ting Gong acquired the data; Zheng Ao, Zhimin Wu and Yiyu Zhang analyzed and interpreted the data; Qun Hu and Linjun Hong contributed to contributed materials; Zheng Ao and Zhimin Wu wrote and revised the paper.

## Supporting information

**S1 Table. Summary of mapping statistics in LW and MW placenta**

**S1 Fig. Volcano map of differential expressed genes between LW and MW placentas**.

**S2 Fig. Transfection efficiency of RACK1 overexpression vector determined by qPCR**.

**P < 0.001.

